# Repurposing The Dark Genome. I - Antisense Proteins

**DOI:** 10.1101/2023.03.15.532699

**Authors:** Mohit Garg, Pawan K. Dhar

**Author notes:** corresponding author Pawan K. Dhar, Ph.D., Professor & Head, Synthetic Biology Group, School of Biotechnology, Jawaharlal Nehru University New Delhi 110067.

## Abstract

From the functional standpoint, the genome may be considered a collection of three types of sequences: protein encoding, RNA encoding, and non-expressing. Based on the standard sequencing and annotation work, it is now well accepted that a small proportion of the genome has been allocated the job of encoding proteins, while most of the genome encodes RNA, and some DNA sequences are not used for expression. The exact ratio among these three types of sequences varies based on the organism. We asked: Is it possible to artificially encode protein and peptide sequences from naturally non-expressing (dark genome) sequences? This led to proof of the concept of making functional proteins from the intergenic sequences of E.coli (Dhar et al 2009). This study is an extension of the original concept and has been organized around antisense DNA sequences. The full-length antisense gene equivalents in forward and reverse orientations were computationally studied for their structural, cellular location, and functional properties, leading to many interesting observations. The current study points to a huge untapped genomic space that needs to be examined from cell physiology, evolutionary, and application perspectives.

## 1. Introduction

Genomes are the fundamental molecular chassis of organisms providing primary molecular codes for running the cell. Genomes come with various sizes and base pair proportions, hosting a unique arrangement of genes and non-genic regions providing the fundamental blueprint for activating the downstream molecular processes. Historically, DNA has been the main focus of several generations of scientists, as it provides intellectual fuel to understand collective molecular events like replication, transcription, translation, pathways, and so on. Following the discovery of protein-coding genes, the regulatory space defined by RNA-coding genes has received increased attention for the last several decades. It is now accepted that the so-called ‘junk’ is a huge repository of RNA coding genes. Interestingly, the non-expressing part of the genome has only met a computational treatment leading to an unclear understanding of its role in the cell.

To fill up this gap, our lab provided preliminary experimental evidence of artificially encoding intergenic sequences into functional molecules (Dhar et al 2009). Extending the search further, the current study was organized to understand the potential of antisense strands in generating novel biomolecules.

## 2. Material and Methods

The *Escherichia coli* (Strain: K-12 MG1655) genome database and *Saccharomyces cerevisiae* (S288C) genome database and *Drosophila melanogaster* (fruit fly) genome database was used in this study (NCBI Resource Coordinators, 2013). Data was downloaded from the NCBI (National Center for Biotechnology Information) database. 4315 protein-coding genes of *E. coli*, 6017 protein-coding genes of *S. cerevisiae*, and 13,962 protein-coding genes of *Drosophila* were used to generate antisense DNA sequence equivalents.

### 2.1 Translation of Antisense and Reverse Antisense Sequences

Sequences on the minus (-) strand of *E. coli* and *S. cerevisiae* genomes were computationally translated into Protein Sequence with the help of the UGENE tool (Okonechnikov et al., 2012). Only those translated sequences used were full-length equivalents, not interrupted by a stop codon (Figure 1). Sequences that showed stop codons within computationally translated antisense strands were discarded. Expasy-Translation Tool (Gasteiger, 2003) was used to quality check the result. In the antisense strand, both directions were used to predict new genes as shown in Fig.1.

**Fig. 1.**
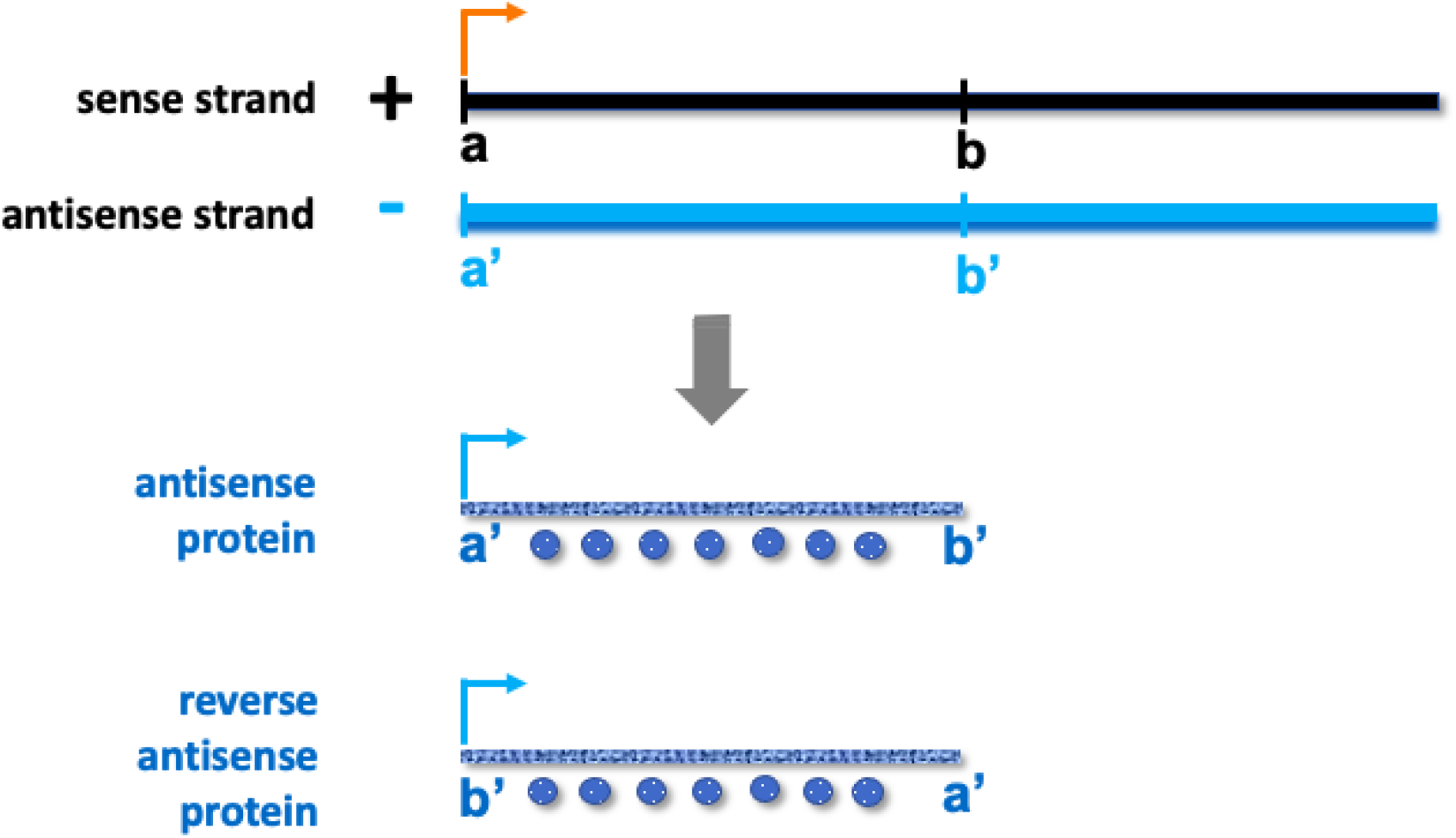
Schematic diagram for making genes and proteins from the antisense strand of DNA (indicated in blue color). Orange colored arrow indicates the direction of natural expression.

### 2.2 Protein Sequence Similarity With Previously Available Database

In the next step, full-length antisense translated sequences were matched against peptides and protein sequences available in the NCBI Non-Redundant Protein Database (Altschul et al., 1990). Only those sequences were included further in the study that did not show any sequence similarity with the available protein sequences.

### 2.3 Physicochemical properties of protein Sequences

Physicochemical properties of Sequences checked by the Expasy-ProtParam tool of SIB Swiss Institute of Bioinformatics (Roy et al., 2011). The ProtParam tool computes various physical and chemical properties of protein sequences such as molecular weight, theoretical pI, instability index, and grand average of hydropathicity (GRAVY) index.

### 2.4 Structure and Function Prediction

Structure and Function were predicted by the I-TASSER Server (Iterative Threading ASSEmbly Refinement) tool (Zhang, 2008). I-TASSER predicts the secondary and tertiary structure of a protein along with probable functions. Subcellular localization of protein was computed by the WoLF PSORT tool (Horton et al., 2007).

### 2.5 Stereochemical properties of protein models

10. The stereochemical properties of the protein model were studied by using Procheck Server. The geometry of amino acids was assessed by the Ramachandran plot. Ramachandran Plot is a two-dimensional (2D) plot between the torsional angles of amino acids phi (φ) and psi (ψ) in protein sequences (Hollingsworth & Karplus, 2010).

## 3. Results

### 3.1 Translation of Antisense and Reverse Antisense Sequences

The computational translation of the antisense DNA sequences showed several full-length gene sequence equivalents (i.e., without stop codon) in the forward and reverse directions (Table 1).

**Table 1.** Full-length gene sequence equivalents in *E. coli, S. cerevisiae, v* and *D. melanogaster*.

| S.No. | Full length protein sequences | *E. coli* (4315 protein-coding genes) | *S. cerevisiae* (6317 protein-coding genes) | *D. melanogaster* (13,962 protein coding gene) |
| --- | --- | --- | --- | --- |
| 1 | antisense proteins | 32 | 10 | 33 |
| 2 | reverse antisense proteins | 208 | 35 | 306 |

### 3.2 Sequence Similarity

The full-length antisense and reverse antisense sequences from *E. coli*, S. cerevisiae, and Drosophila were matched against the existing NCBI protein sequence database. Some sequences are not found to have any similarity with previously available protein databases (Novel protein) and the remaining sequences show similarity in different percentages. (Table 2)

**Table 2.** Sequence similarity of the full-length antisense and reverse antisense sequences of *E. coli, S. cerevisiae*, and *D. melanogaster* (with the existing protein sequence data)

| S. No. | Similarity Range → | Unique<br>(No Similarity ) | Similarity<br>< 30% | Similarity<br>30% - 80% | Similarity<br>above 80% |
| --- | --- | --- | --- | --- | --- |
|  | Full-length sequences |  |  |  |  |
| **1.** | ***E. coli*** |  |  |  |  |
| a. | antisense protein Sequences | 9 | 0 | 18 | 5 |
| b. | reverse antisense protein sequences | 6 | 5 | 52 | 145 |
| **2.** | ***S. cerevisiae*** |  |  |  |  |
| a. | antisense protein sequences | 7 | 0 | 3 | 0 |
| b. | reverse antisense protein sequences | 18 | 1 | 6 | 10 |
| **3.** | ***D. melanogaster*** |  |  |  |  |
| a. | antisense protein sequences | 25 | 1 | 7 | 0 |
| b. | reverse antisense protein sequences | 209 | 32 | 56 | 9 |

### 3.3 Physicochemical properties

Antisense and reverse antisense encoded proteins were studied for their physicochemical properties (Table 4). The molecular Weight of antisense proteins is found to range from 3.8 KDa to 43.0 KDa, with Isoelectric Point (pI) value ranging from 3.80 to 13.04 (pI <7 show acidic nature whereas pI >7 indicate basic nature), Instability index ranged from 4.5 - 134, hydropathicity value (GRAVY) ranged from - 0.496 to 0.672 showing interaction with water (low GRAVY score indicates better interaction with water), with G-C ratio ranging from 33.3 - 64.4.

**Table 3.**
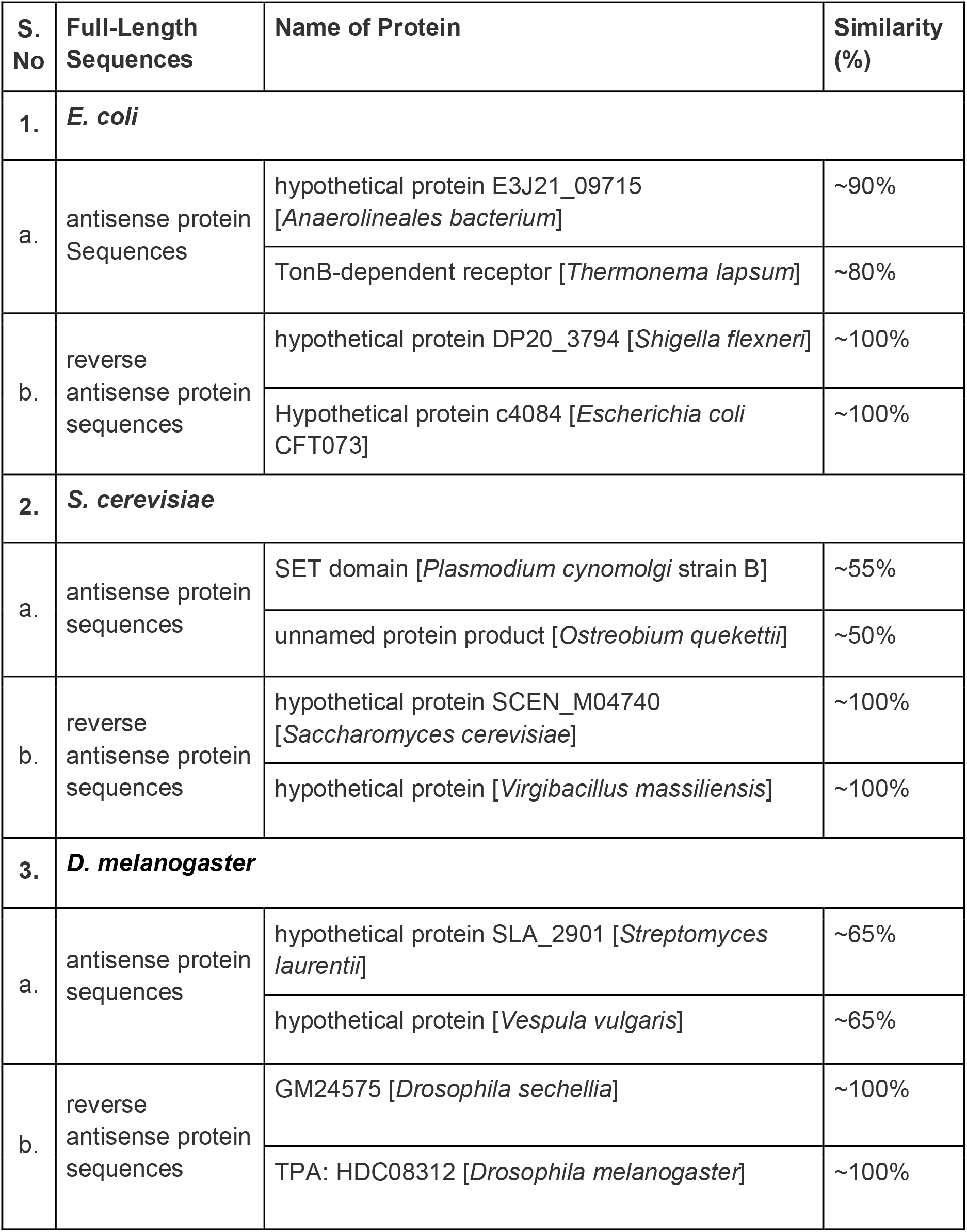
Predicted identification and similarity of putative antisense and reverse antisense proteins of *E. coli, S. cerevisiae*, and *D. melanogaster*.

**Table 4.** Physicochemical properties of hypothetical antisense and reverse antisense proteins from *E. coli, S. cerevisiae, and D. melanogaster*.

| Type | amino acid residues Length (range) | Molecular weight (kDa) (range) | Theoretical pl (range) | Instability index (range) | Grand average of hydropathicity (GRAVY) (range) | G-C % (range) |
| --- | --- | --- | --- | --- | --- | --- |
| ***E. coli*** |  |  |  |  |  |  |
| Antisense Protein | 31 - 78 | 3.8-9.1 | 6.69-12.55 | 10 - 118 | -0.496 - 0.672 | 40.9-50.8 |
| Reverse Antisense Protein | 44 - 70 | 5.1-8.3 | 6.68-11.84 | 30 - 71 | -1.792 - 0.468 | 39.5-59.7 |
| ***S. cerevisiae*** |  |  |  |  |  |  |
| Antisense Protein | 30 - 94 | 3.8-11.7 | 5.97-12.16 | 33 - 86 | -0.977 - 0.472 | 33.3-47.3 |
| Reverse Antisense Protein | 32 - 121 | 4.0-13.3 | 4.43-11.73 | 10 - 67 | -1.322 - 0.866 | 31.7-53.4 |
| ***D. melanogaster*** |  |  |  |  |  |  |
| Antisense Protein | 33 - 178 | 3.9-20.6 | 7.75-12.02 | 11 - 94 | -0.963 - 1.418 | 44.4-64.4 |
| Reverse Antisense Protein | 32 - 400 | 3.6-43.0 | 3.88-13.04 | 4.5-134 | -1.287 - 1.586 | 34.3-64.3 |

### 3.4 Structure and Function Prediction

Structure and function prediction of dark proteins predicted by I-Tasser showed a favorable C-score i.e., within the range of -5 to +2.

The secondary and tertiary prediction of protein structures (predicted by I-TASSER) along with normalized B factor (BFP) profile were found within a favorable range. BFP shows experimental structure stability of proteins. Subcellular localization (WoLF PSROT server) showed predominant localization of proteins to mitochondria and several cytoplasmic destinations, while some were found to be secretory proteins (Table 5).

**Table 5.** Predicted structure and function prediction of putative antisense and reverse antisense proteins of *E. coli, S. cerevisiae*, and *D. melanogaster*.

| Organisms | structure prediction (C-score range) | function prediction (types) | subcellular localizations |
| --- | --- | --- | --- |
| *E. coli* | -1.76 to -3.29 | transferase, electron transport, oxidoreductase, hydrolase | secreted, mitochondrial, cytoplasmic, and nuclear |
| *S. cerevisiae* | -1.33 to -4.02 | transferase, hydrolase, electron transport, protein transport, oxidoreductase | secreted, mitochondrial, cytoplasmic, and nuclear |
| *D. melanogaster* | -0.67 to -4.11 | hydrolase, lyase, ligase, oxidoreductase, isomerase, Transferase, | secreted, mitochondrial, peroxisomal, nuclear, and cytoplasmic |

### 3.5 Stereochemical properties

Ramachandran Plot was used to study the stereochemical properties of proteins (Fig 4). Most of the residues were found in favorable regions indicating a good protein folding profile.

**Fig. 2.**
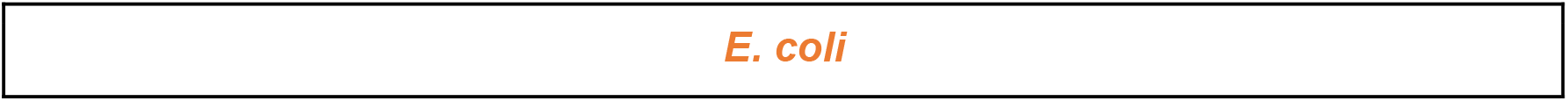
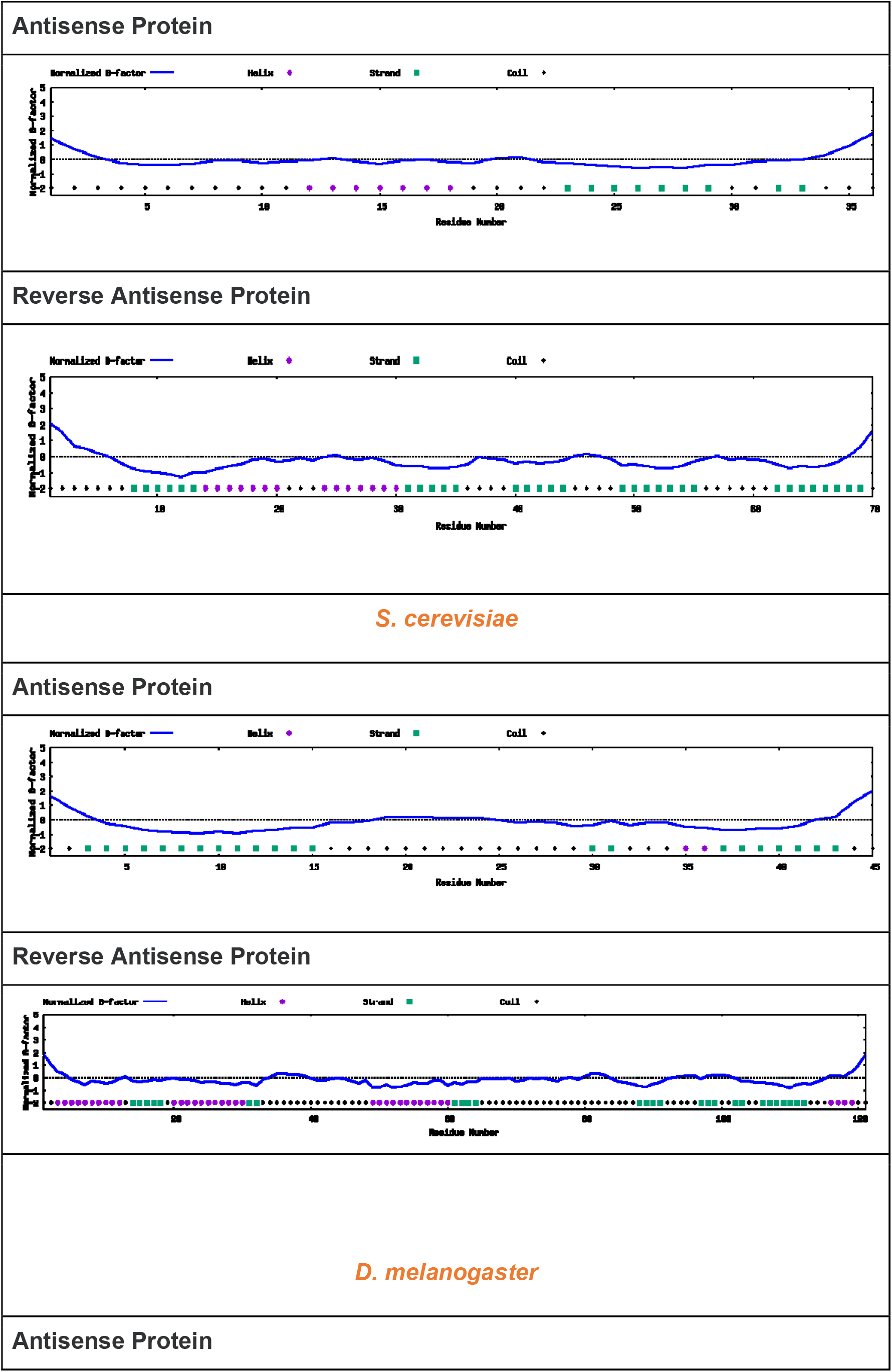
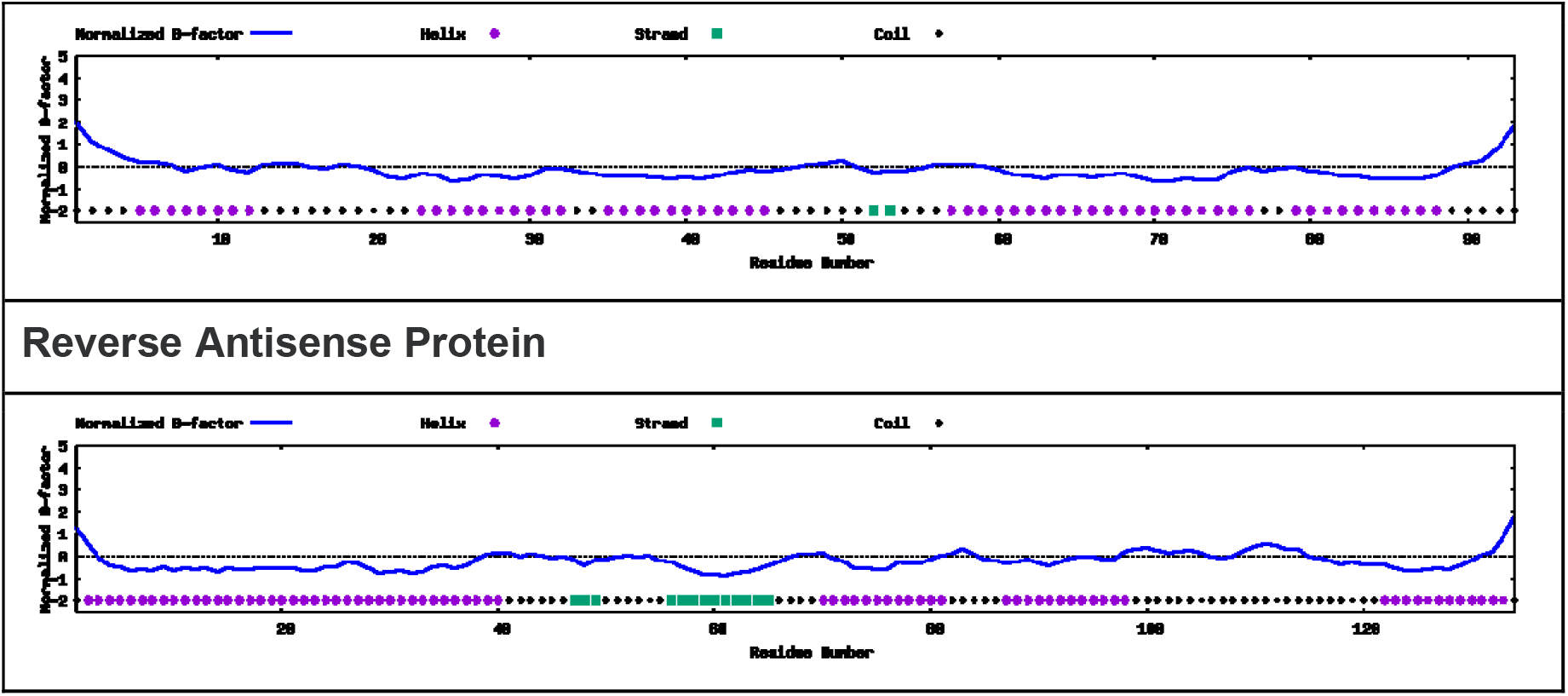
Normalized B-Factor Profile of antisense and reverse antisense proteins in E.coli, S.cerevisiae, and D.melanogaster with most of the residues below 0 indicating the stability of proteins.

**Fig. 3.**
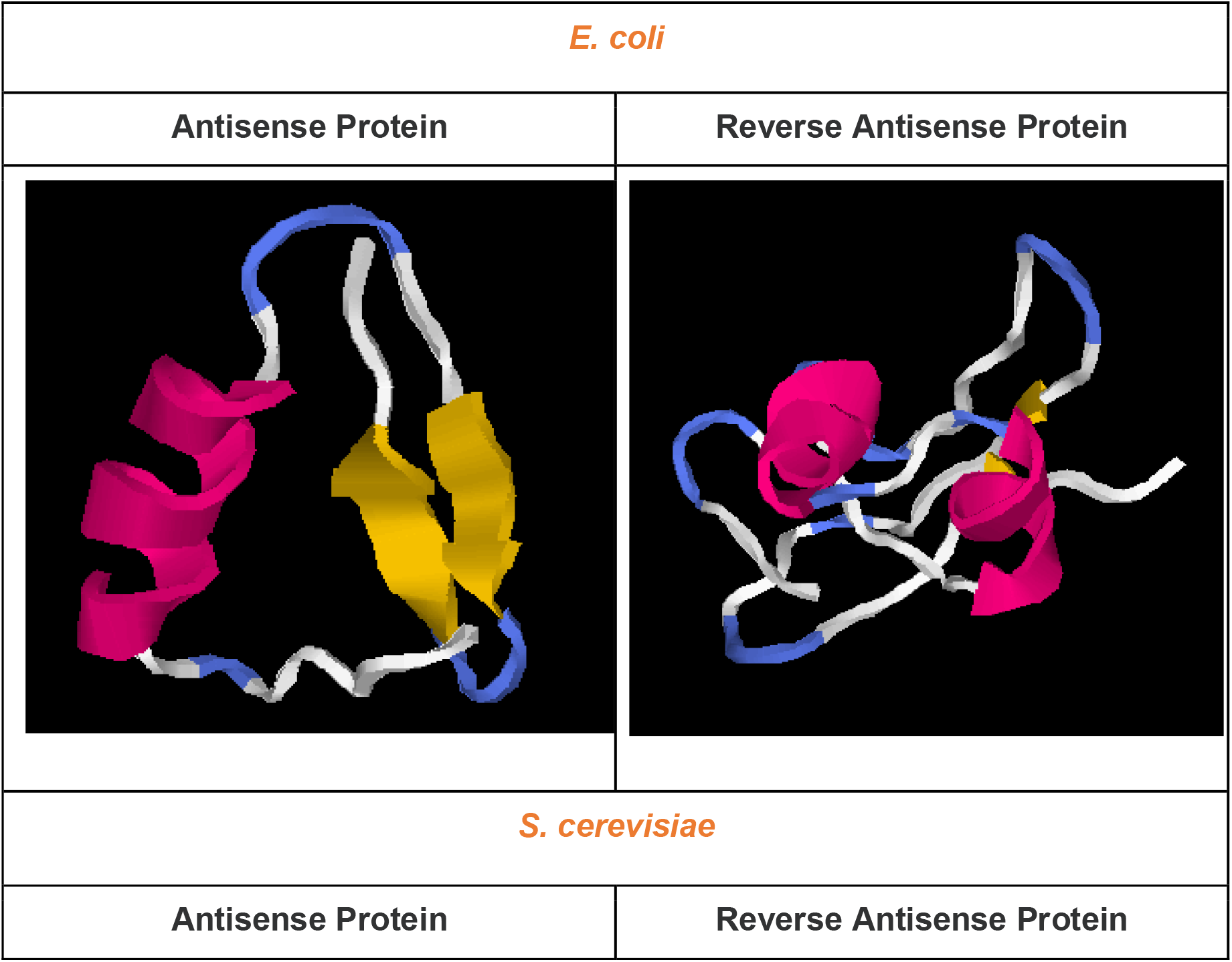
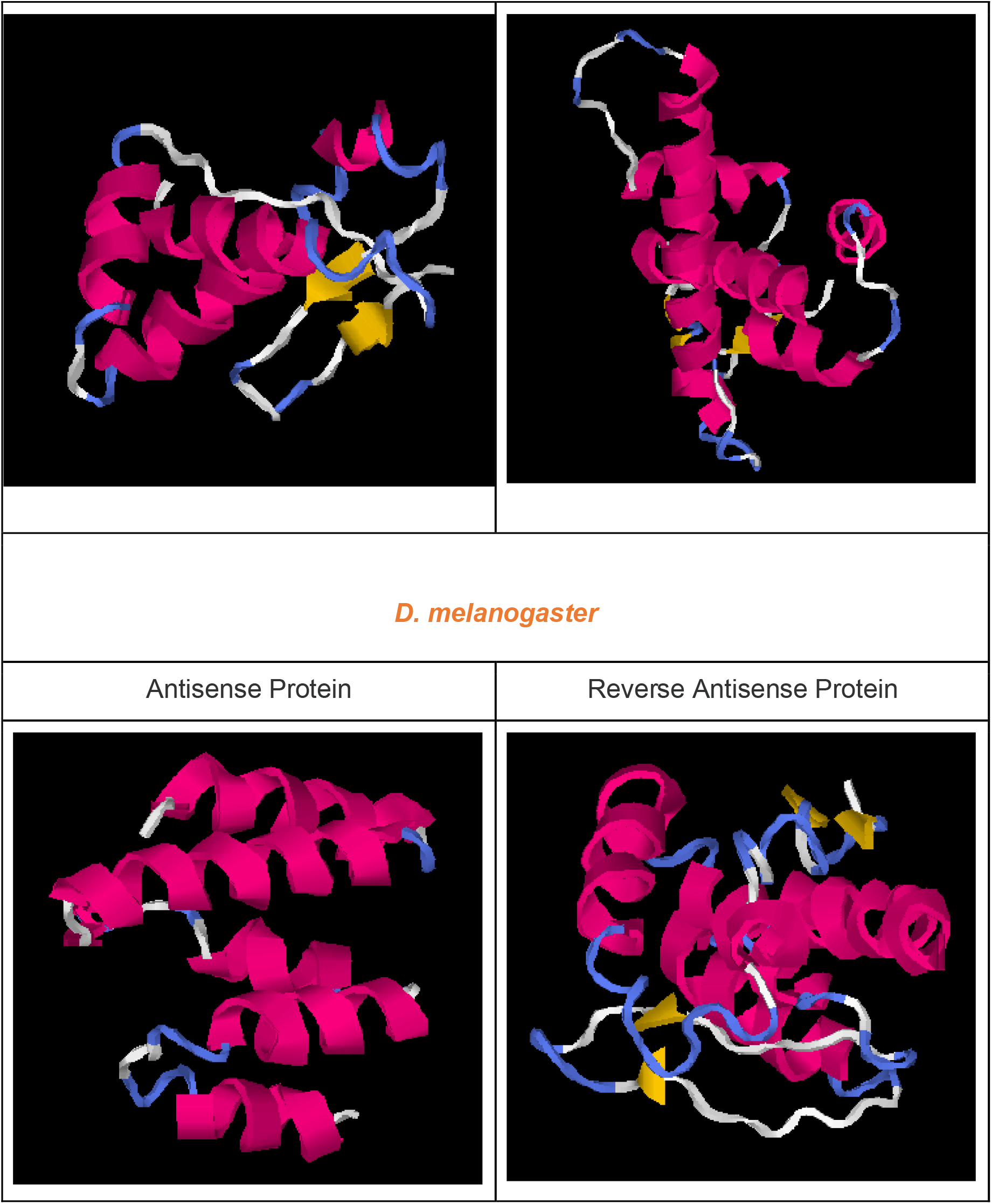
Tertiary structure of antisense and reverse antisense protein sequences sampled from E.oli, S.cerevisiae, and D.melanogaster resembling naturally expressing proteins (prediction tool: I-TASSER)

**Fig. 4.**
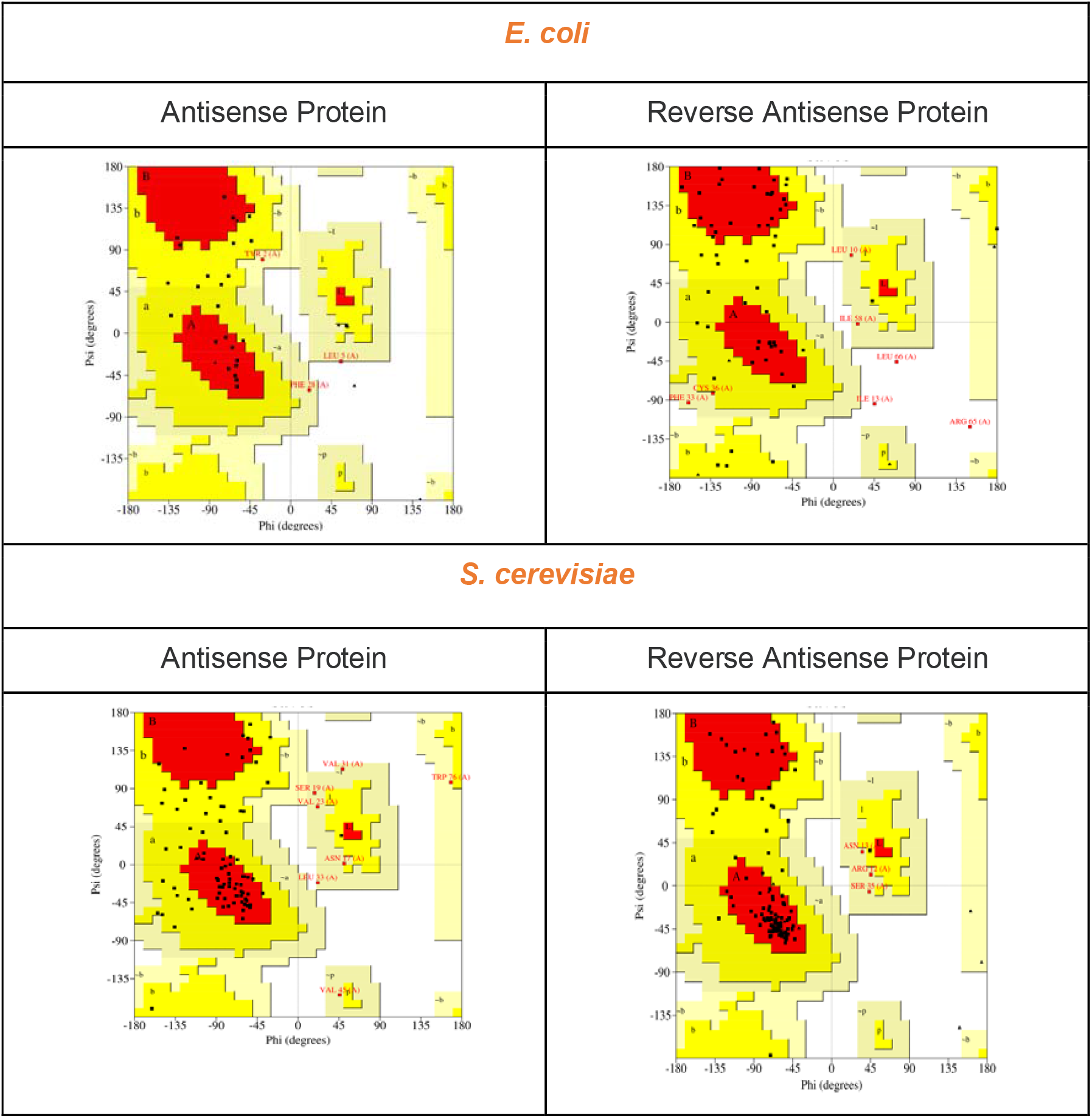
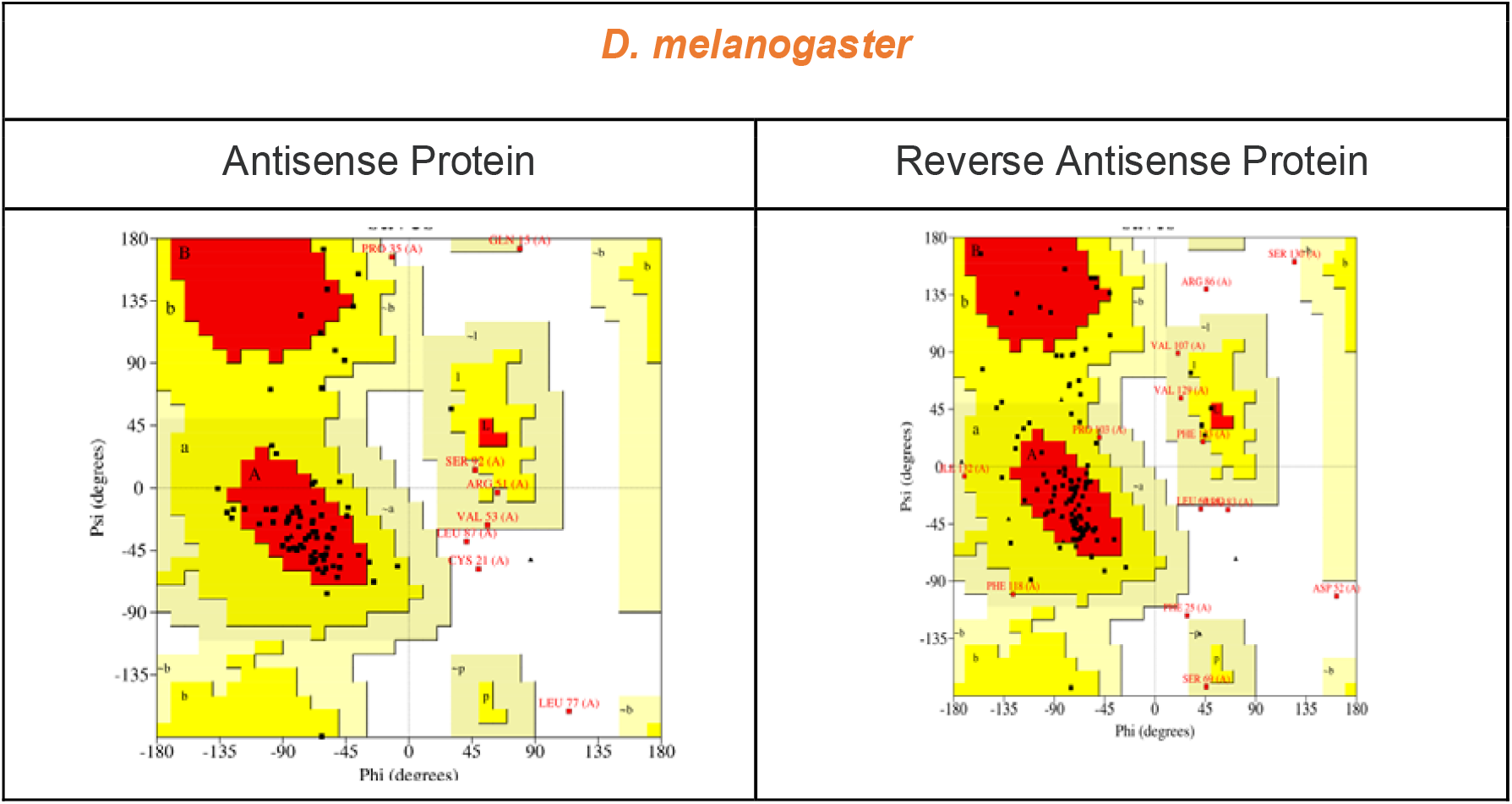
Ramachandran Plot of antisense and reverse antisense protein sequences of E.coli, S.cerevisiae, and D.melanogaster. Most of the amino acid residues were found to be present in red and dark yellow regions.

## 4 Discussion

The term ‘dark-genome’ refers to fully dark (never expressing), partially dark (RNA expressing), and virtually dark (pseudogenes) sequences that can be artificially expressed into functional peptides and proteins. The fully dark sequences comprise antisense, reverse coding, highly repetitive sequences, and intergenic sequences. The “partially dark” genome comprises sequences that didn’t go beyond transcription i.e., tRNA, long ncRNA, ribosomal RNA, and intron sequences. The “virtually dark” genome comprises sequences like pseudogenes that were previously expressed but were retired during evolution.

This study using *E*.*coli, S*.*cerevisiae*, and *D*.*melanogaster* is an extension of our previous work (Dhar et al., 2009) where intergenic sequences of E.coli were artificially expressed into cell growth inhibiting proteins. Subsequently, it was observed that pseudogenes also have the potential to generate stable and functional proteins, if expressed (Shidhi et al., 2015)

Given that genes are found on both strands, the term antisense does not indicate an “end-to-end” stretch of complementary DNA. Thus, sense and antisense only indicate a specific arrangement on a particular genomic address. In this study, we considered full-length hypothetical genes on the antisense strands in the forward and reverse directions. DNA sequences that showed stop codons on computational translation were discarded.

Strong evidence of unutilized genomic potential in the form of hypothetical full-length antisense proteins and reverse antisense proteins were observed in *E*.*coli* (0.7% and 5.1%), *S*.*cerevisiae* (0.15%, 0.5%) and *D*.*melanogaster* (0.2%, 2.1%), respectively. The physicochemical properties of these molecules favored the possibility of a well-defined structure and function.

Most of the proteins considered in this study were predicted to be located in various cellular locations with some proteins showing a possibility of secretory nature. The functional prediction tools mapped many of these proteins to transporter and enzyme functions.

To our best knowledge, this is the first report that describes the repurposing of antisense DNA sequences toward generating functional biomolecules. An additional parameter of “reverse antisense” proteins was considered in this study, to raise more evolutionary questions and increase the inventory of synthetic biomolecules. In the future, more work is needed to release the raw antisense and reverse antisense data in the form of a publicly available database and experimentally validate some of the key predictions.

## Author Contributions

PKD conceived the idea of making functional proteins from antisense DNA strands, supervised the work and wrote final version of the manuscript. MG performed all the computational work, and drafted the first version of the manuscript.

## Acknowledgment

MG would like to express its warmest thanks to Mr. Shubham Garg for helping in some of the key computational work and CSIR for providing NET-JRF Scholarship. PKD expresses sincere thanks to JNU for the critical lab support and to Prof. Binay Panda (JNU) for extremely helpful discussions.

## Notes

### Competing Interest Statement

The authors have declared no competing interest.

### Summary of Updates

1. Author contributions 2. Typos in the previous version

